# Toxicity Prediction Tools Do Not Generalize to Novel Chemistry

**DOI:** 10.64898/2026.10.26.761698

**Authors:** Martin Weiss, Circe Hsu, Cailum M. K. Stienstra, Amy Caudy, Antonio H. de O. Fonseca

## Abstract

Computational toxicity prediction rests on an assumption that is rarely tested directly: that molecular structure carries enough information to anticipate whether a compound will have adverse effects in human patients. We benchmark toxicity prediction methods on a curated panel of 569 approved drugs, labeled toxic if they carry a black-box warning or were withdrawn from the market. We evaluate 35 model readouts spanning fingerprint-based QSAR, fine-tuned foundation encoders, and toxicity-targeted tools. No learned representation outperforms a plain fingerprint, suggesting the bottleneck is not the sophistication of the representation. Of the tools trained specifically for toxicity prediction, only one performs above chance, and none matches what approval year alone explains, a predictor containing no chemistry at all. No tool dedicated to drug-induced liver injury (DILI) reliably beats chance on our panel, and the external set where some succeed is one that approval year alone largely predicts. These results point to a limit of molecular structure as an input, and suggest that progress will depend on inputs that capture exposure and metabolism rather than better models over structure alone.

## 1 Introduction

Predicting human clinical toxicity from molecular structure is a central promise of machine learning for drug discovery. Various methods exist to model toxicity from a chemical input: classical Quantitative Structure–Activity Relationship (QSAR) over circular fingerprints, message-passing graph networks, pretrained chemical language models, 3D structure encoders, and instruction-tuned therapeutic models [Rogers and Hahn, 2010, Yang et al., 2019, Ross et al., 2022, Zhou et al., 2023, Wang et al., 2025]. These models achieve strong benchmark performance, suggesting that they are suitable for use in drug development pipelines. However, benchmark performance is usually measured under random or scaffold splits, leaving test compounds close to the training set. Such benchmarks fail to measure how models will perform when evaluated on novel structures, such as a new drug class. This gap between reported and prospective accuracy is a recurring theme in critical reviews of the field [Bender and Cortes-Ciriano, 2021]. We therefore ask a simple question: *Can a model predict, from structure alone, whether a novel drug is clinically toxic in humans?* We assembled a benchmark of 569 drugs from ChEMBL 37 [Zdrazil et al., 2024] consisting of recent approvals, withdrawals, and black-box warnings, all absent from invitroDB with sub-0.7 ECFP4 (extended-connectivity fingerprint of radius 2) Tanimoto similarity to the 8,789 reference compounds in invitroDB, and evaluated the models of interest on this cleaned benchmark set.

## Contributions

- **A novelty-gated clinical toxicity panel**. We assemble a dataset with 569 approved drugs labeled by regulatory outcome, absent from invitroDB and below 0.70 ECFP4 Tanimoto similarity to any compound in it. We then evaluate 35 model readouts under two protocols: models fit on the panel by cross-validation, and models applied frozen as released.
- **Representation sophistication is not the bottleneck**. We test state-of-the-art foundation structure encoder models fit on the panel for toxicity prediction, and show that no learned representation outperforms a raw ECFP4 fingerprint.
- **Tools built for toxicity prediction do not transfer to novel chemistry**. We apply tools trained specifically for toxicity prediction to our novel drug panel, and show that no DILI-specific tool clears chance on it.

## 2 Related work

### Structure-to-Property and Surrogate Models

Toxicity prediction maps a structure to a toxicity score, and models differ mainly in their approach to structural featurization. Popular models utilize fixed extended-connectivity fingerprints (ECFP) feeding a classifier [Rogers and Hahn, 2010], an approach still used in regulatory QSAR systems in compliance settings, while ProTox [Banerjee et al., 2024] combines molecular similarity with fragment propensities. Message-passing networks [Yang et al., 2019, Heid et al., 2024], graph-attention models [Xiong et al., 2021] and self-supervised encoders [Chithrananda et al., 2020, Ross et al., 2022, Zhou et al., 2023] learn structural features instead. A second group supervises on biological activity rather than clinical outcome, through regression on assay panels [Mayr et al., 2016, Swanson et al., 2024], proxy labels as features [Seal et al., 2024], or prediction of the full bioactivity profile [Arturi et al., 2025]. Using such a profile as a claim about clinical harm requires further mapping, from assay bioactivity to toxicity in a treated organism, and that mapping remains unresolved. Linking the two on a dose basis needs physiologically-based toxicokinetic modelling, and even with those models in place, the best assumption set outperformed a random parameterization in 71% of in vitro to in vivo endpoint comparisons rather than in nearly all of them [Honda et al., 2019]. Used directly instead, assay scores have proved fragile: one reanalysis of rodent carcinogenicity prediction found accuracy no better than chance for roughly three-quarters of the chemicals [Cox et al., 2016]. The most recent entrant, TxGemma, covers 66 therapeutic development tasks with a single model and no per-task head [Wang et al., 2025], which is appealing where individual toxicity labels are scarce. Its pretraining corpus is undisclosed, so how much of a given benchmark it has already seen cannot be established, only bounded, which is why it is the only entry in our audit reported as a lower bound rather than a measurement.

### Evaluation and Leakage

These families differ in representation and supervision. The representation ranges from a fixed descriptor to a learned one, and the supervision from the target label to surrogate assays to no toxicity labels at all. Two properties are common to all of them. None represents the metabolic transformation of the administered molecule, which is where much toxicity originates. In addition, each reports its accuracy on benchmarks whose test compounds resemble its own training set: random splits leave near-duplicate analogs on both sides [Wu et al., 2018, Sheridan, 2013], ligand-based benchmarks are separable by analog bias alone [Wallach and Heifets, 2018], and performance is worse under scaffold splits and on activity cliffs [Deng et al., 2023, van Tilborg et al., 2022] (Table 1 defines these terms). Those endpoints are almost all biochemical, however, and where a toxicity label appears, as with Tox21 in MoleculeNet, it is an in vitro assay panel rather than a clinical outcome. Whether these findings carry over to a clinical toxicity label, and whether the relevant unit of leakage is the split or each tool’s own training corpus, both remain open, and we investigate both in our work.

**Table 1:** Glossary. Terms used throughout the paper.

| Term | Definition |
| --- | --- |
| Activity cliff | A pair of near-identical structures with very different activity. |
| AMES | The Ames test, a bacterial assay for whether a compound causes DNA mutations. |
| Analog bias | Benchmark separability that comes from which chemical series a compound belongs to, rather than from the property being predicted. |
| Black-box warning | The FDA’s strongest labeling requirement, applied for serious or life-threatening adverse effects. |
| ClinTox | A benchmark that labels drugs by whether they failed clinical trials for toxicity. |
| DILI | Drug-induced liver injury; a leading cause of post-approval withdrawal. |
| ECFP4 | Fixed binary fingerprint marking which circular atom environments, out to radius 2, a molecule contains. |
| Endpoint | One specific predicted outcome (hERG block, Ames mutagenicity, skin sensitization), not overall clinical harm. |
| Frozen | No parameter fit on the panel. For the structure encoders (Class 2), the encoder weights only, since a linear probe is still fit on panel labels. |
| GIN, Grover | Graph neural networks that learn from a molecule’s atoms and bonds. GIN is the graph isomorphism network, and Grover is a graph transformer pretrained on unlabeled molecules. |
| Grade A | LiverTox’s top likelihood category, a well-known cause of liver injury with at least 50 published cases. |
| hERG | A cardiac potassium channel whose block can disturb heart rhythm and cause arrhythmia. |
| invitroDB | The public data release of ToxCast, the EPA’s in vitro screening program, with 8,789 compounds across hundreds of assays. |
| Leakage | Test compounds present in training, whether through the split or through a tool’s own corpus. |
| LiverTox | NIH reference grading a drug’s hepatotoxicity from the published case literature, on a likelihood scale from A to E. |
| Panel | Our 569-drug evaluation set. |
| QSAR | Quantitative structure–activity relationship, a model mapping structural features to an activity or toxicity label. |
| Readout | One scored combination of model, classification head and endpoint; a single tool can yield several. |
| Scaffold split | A train–test split that puts molecules with different core ring systems on opposite sides, so test molecules differ structurally from training molecules. |
| Tanimoto similarity | Fraction of fingerprint bits shared between two molecules. |
| ToxCast | EPA program screening chemicals across in vitro assays to prioritize them for toxicity testing. |
| Toxic / clean | Our labels. Toxic means a black-box warning or withdrawal, and clean means approval with neither. |

## 3 Methods

### Datasets

We evaluate 35 model readouts on 569 approved drugs drawn from ChEMBL 37 [Zdrazil et al., 2024], labeled *toxic* if a drug carries a black-box warning or was withdrawn (235 compounds, 148 of which carry an explicit ChEMBL toxicity annotation, and others are identified only by the black-box warning or withdrawal) and *clean* if approved with neither (334). Every compound is absent from invitroDB [Dix et al., 2007] and below 0.70 maximum ECFP4 Tanimoto similarity to all 8,789 of its reference compounds. The cutoff is applied to the panel and not to individual models, and Appendix A.2 examines how results change with similarity to invitroDB. Because that criterion is defined against invitroDB rather than any individual tool’s corpus, we also verify each tool’s training set for panel overlap, matching on 2D InChIKey joined with canonical SMILES. As an external dataset, we use the 314-compound evaluation set of Bergen et al. [2025] (hereafter referred to as LiverTox-314), which consists of 45 *DILI-positive* and 269 *DILI-negative* compounds labeled by clinical experts.

### Models and Evaluation Setups

The roster spans four Classes, grouped by model family and by how much each model can learn from the panel (Table 2, Appendix A). Throughout, *frozen* means no pretrained weight is updated. Class 0 sets the floors, all fit with 5-fold cross-validation on the panel: fixed-representation fingerprint and descriptor models, plus two metadata probes carrying no chemistry at all (approval year and therapeutic class). Class 0 also holds five readouts of a pipeline built to predict bioactivity across 471 invitroDB assays from molecular structure. Two of those readouts fit an aggregating head on panel labels, drawn with open markers in Figure 1; the other three are frozen. Class 0 establishes the level a learned representation (Class 2) must clear on the same folds and the same supervision. Class 1 holds published toxicity prediction tools, evaluated frozen and used exactly as released, which is how a drug-development team would use them. Class 2 holds pretrained molecular foundation encoders, finetuned on the panel and, where a checkpoint exists, also run with the encoder frozen and a linear probe fit over its embeddings. Class 3 takes a fingerprint-plus-descriptor model trained on animal-study data and applies it to our human panel.

**Table 2:** Model roster by Class and family. Panel CV readouts are fit on the panel by cross-validation. This includes the linear probes over the molecular encoders, whose encoder weights are held fixed but whose classification head is fit on panel labels, so they are marked *frozen* in Figure 1 in that sense only. Frozen readouts in the column below are trained entirely off-panel and applied without any fitting to it. All 35 readouts appear individually in Figure 1; Appendix A.1 describes each Class.

| Class | Model family | $n$ | | Readouts |
| --- | --- | --- | --- | --- |
|  |  | CV | frozen |  |
| 0 | ECFP4 baseline | 8 | — | raw ECFP4 (LR, XGB, RF); ECFP4 + physchem (LR, XGB); physchem; approval year; therapeutic class |
| 0 | Bioactivity pipeline | 2 | 3 | endpoint pattern (RF, LR) fit on panel; endpoint pattern, summary score and in vivo bridge applied frozen |
| 1 | Toxicity prediction | — | 12 | seven ADMET-AI heads; two Tox21-lineage readouts; DILIPredictor; DILIGeNN; TxGemma |
| 2 | Structure encoders | 7 | — | Uni-Mol, MolFormer, ChemBERTa (finetuned and linear probe); Chemprop (finetuned only) |
| 3 | In vivo | — | 3 | ToxRefDB ECFP4 + physchem (LR, XGB, RF) |
| Total |  | 17 | 18 | 35 readouts |

**Figure 1:**
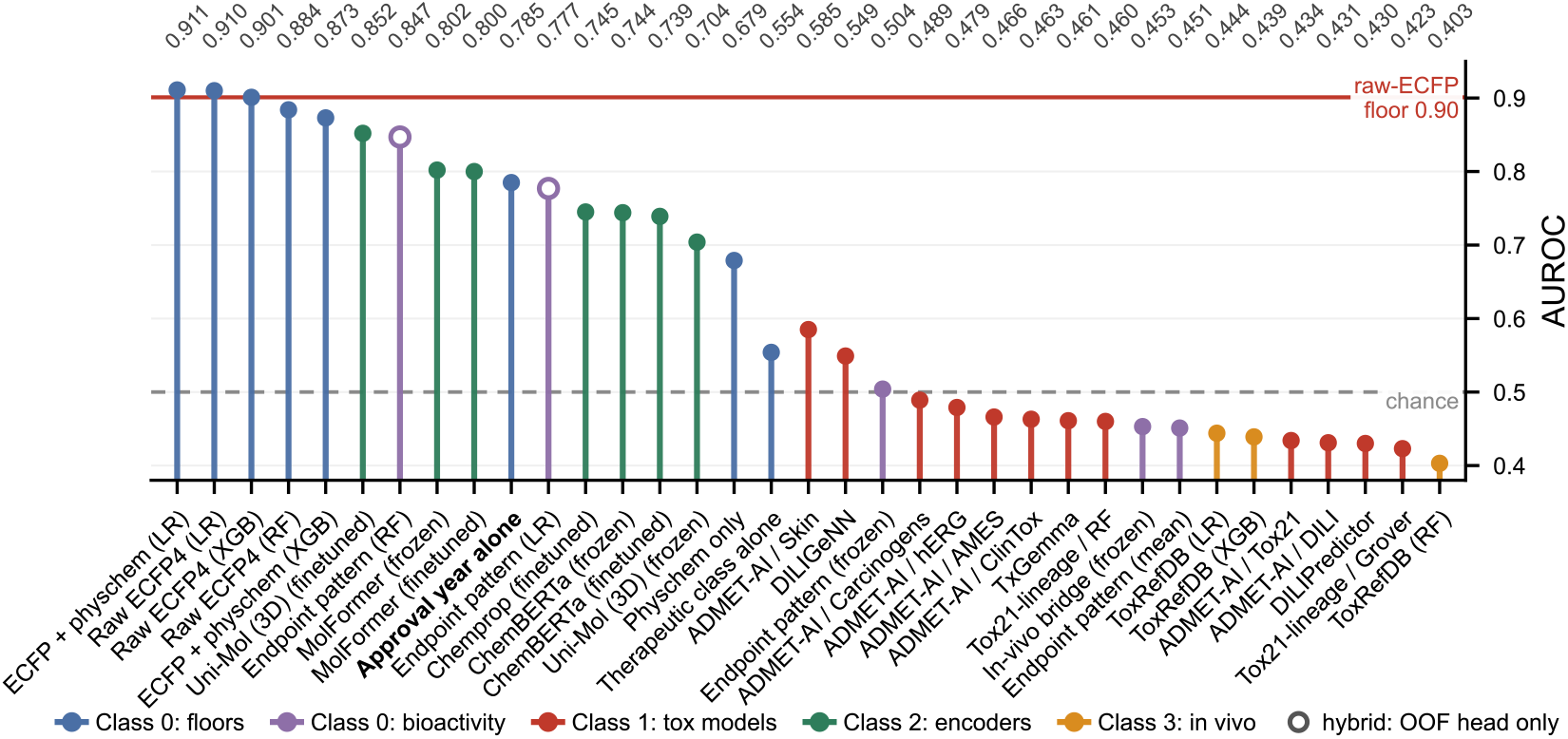
Model performance. AUROC for all 35 models on the 569-drug panel. Classes follow Table 2. Open markers are the hybrid setup: endpoint predictions from an off-panel model, aggregating head fit on panel labels. For Class-2 encoders, *frozen* denotes a linear probe over fixed embeddings, head still fit on panel labels; elsewhere it denotes no panel fitting at all. Heads in parentheses: logistic regression (LR), random forest (RF), XGBoost (XGB) (see Table 1 and Appendix A for definitions and tool names).

## 4 Results and Discussion

We report the two evaluation setups separately, since a model fit on the panel (Class 0 and Class 2) and a frozen model used for toxicity prediction (Class 1 and Class 3) are not competing on equal terms. First, we compare fixed and learned representations of molecular structure in predicting toxicity.

### Fixed versus learned representations

Among models fit within the 569-drug panel on identical 5-fold splits, raw ECFP4 reaches AUROC 0.910 with logistic regression, 0.901 with XGBoost and 0.884 with random forest, and no learned representation (Class 2) matches it (Figure 1). The strongest is Uni-Mol’s 3D encoder at 0.852, followed by MolFormer at 0.800, Chemprop at 0.745 and ChemBERTa at 0.739. Concatenating physicochemical descriptors with fingerprints yields 0.911 versus 0.910, so those descriptors add nothing beyond the fingerprint. The frozen-probe comparison locates where the encoders’ performance originates: for both chemical language models a frozen linear head matches full finetuning, at 0.802 against 0.800 for MolFormer and 0.744 against 0.739 for ChemBERTa, so the pretrained representation is already close to linearly separable for this label and end-to-end adaptation contributes little at *n* = 569. Only Uni-Mol gains from adaptation, its frozen embedding reaching 0.704. Regression on approval year achieves 0.785, above three of the four encoders. Among the frozen tools (Class 1 and 3) on the toxicity label, most fall below chance, and only the ADMET-AI skin-sensitization head has a CI excluding it (0.585, CI 0.536–0.631); the next best, DILIGeNN, does not (0.549, CI 0.499–0.599). However, both perform worse than approval year alone (0.785), which we include as a reference for what a predictor with no chemistry can recover, not as a competing method. The toxic label pools black-box warnings and withdrawals, but restricting positives to the 148 drugs with an explicit toxicity annotation in ChEMBL changes no tool’s AUROC by more than 0.025 (Table 4, Appendix A). Relaxing the similarity cutoff does not change these conclusions (Appendix A.2).

**Table 3:**
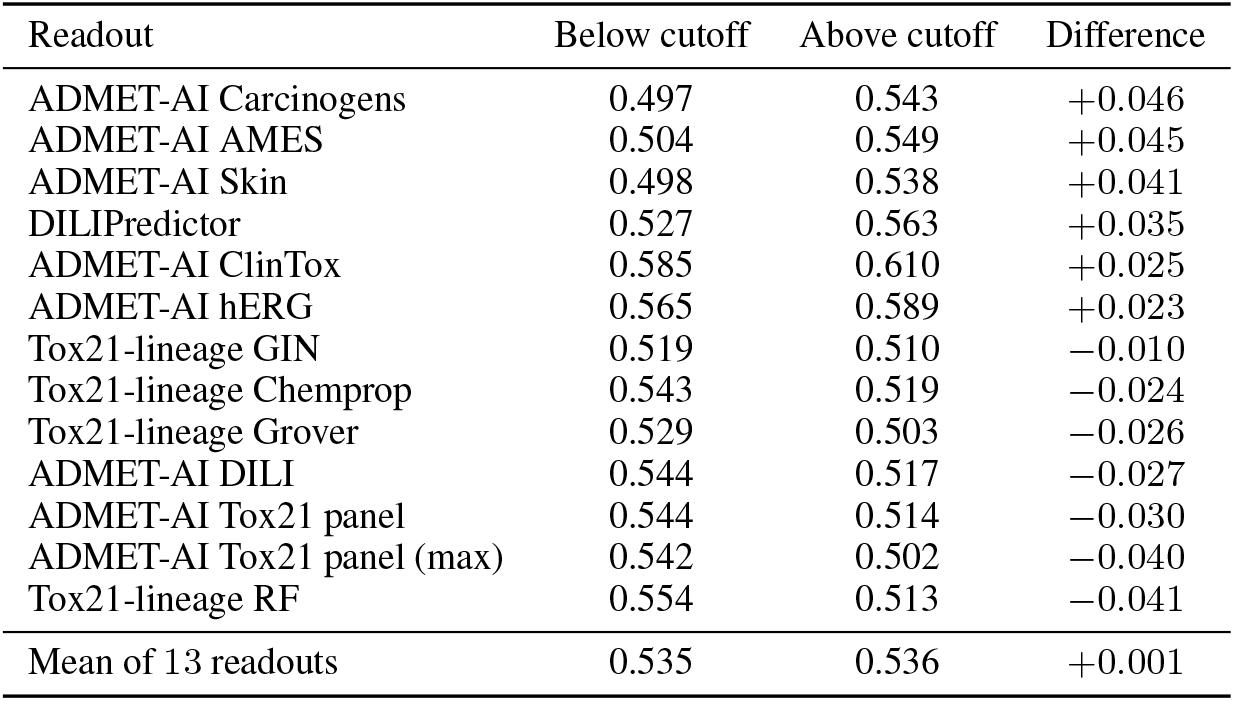
Frozen-tool AUROC below and above the novelty cutoff. For each of the 13 frozen readouts, the mean AUROC across similarity bins to invitroDB below 0.70 and at or above 0.70, weighted by the number of drugs per bin, on a cohort of 1,945 approved drugs selected with no similarity or first-approval cutoff. Weighted means average the bin AUROCs and are not pooled AUROCs. Six readouts score higher above the cutoff and seven score lower, with a median difference of *−*0.010.

**Table 4:** Frozen toxicity tools on organ-toxicity positives. AUROC for the 12 Class-1 tools, ordered by organ-tox AUROC. *Broad label* scores all 235 toxic drugs against the 334 clean drugs; *Organ-tox only* keeps as positives the 148 toxic drugs with an explicit ChEMBL toxicity annotation. Values follow removal of structures matching each tool’s training set on 2D InChIKey or canonical SMILES, so *Positives* is the number of organ-toxicity drugs remaining per tool minus that tool’s matches. CIs are 95% bootstrap on the organ-tox-only AUROC; Δ is organ-tox only minus broad. Narrowing the label moves AUROC by *−*0.013 to +0.025, well inside the CIs, so the broad black-box and withdrawal label does not explain the panel results.

| Tool | Broad label | Organ-tox only | 95% CI | $\Delta$ | Positives |
| --- | --- | --- | --- | --- | --- |
| ADMET-AI / Skin | 0.586 | 0.591 | 0.535–0.645 | +0.005 | 148 |
| DILIGeNN | 0.549 | 0.557 | 0.492–0.615 | +0.008 | 109 |
| ADMET-AI / Carcinogens | 0.491 | 0.514 | 0.455–0.573 | +0.023 | 146 |
| ADMET-AI / hERG | 0.477 | 0.489 | 0.429–0.547 | +0.012 | 138 |
| TxGemma | 0.500 | 0.487 | 0.408–0.561 | −0.013 | 89 |
| ADMET-AI / ClinTox | 0.484 | 0.486 | 0.421–0.554 | +0.002 | 106 |
| DILIPredictor | 0.445 | 0.470 | 0.408–0.537 | +0.025 | 117 |
| ADMET-AI / AMES | 0.462 | 0.470 | 0.410–0.528 | +0.008 | 139 |
| Tox21-lineage / RF | 0.470 | 0.467 | 0.409–0.530 | −0.003 | 133 |
| ADMET-AI / Tox21 | 0.451 | 0.464 | 0.405–0.524 | +0.013 | 133 |
| Tox21-lineage / Grover | 0.434 | 0.448 | 0.385–0.509 | +0.014 | 133 |
| ADMET-AI / DILI | 0.434 | 0.435 | 0.379–0.493 | +0.001 | 133 |

### Drug-induced liver injury

Three of the Class 1 tools are purpose-built for hepatotoxicity, so we assess performance against DILI labels on two sets: the LiverTox-314 and the 569-drug panel scored on DILI (Figure 2). To account for leakage, both panels remove, per tool, the compounds matching their own training set. On LiverTox-314, four of eight readouts exclude chance, while none do on our panel, which carries only 22 DILI-positives and is therefore a consistency check rather than an independent negative result. The two DILI-specific tools that clear chance on the LiverTox-314 DILI label, DILIPredictor (0.601) and DILIGeNN (0.598), do so on a set whose labels approval year alone largely predicts. ADMET-AI Tox21 (0.703) and Tox21-lineage (0.673) show the same pattern. LiverTox-314 grades a drug’s hepatotoxicity from the published case literature, which requires years of post-approval exposure to accumulate. Its hepatotoxic compounds are therefore more skewed towards older compounds (median approval year of 1995, *n* = 33) than our panel (2012, *n* = 13) (Figure 4). Within the LiverTox-314 set, approval year and the label are themselves confounded, with DILI-positives sitting at a median of 1995 against 2017 for DILI-negatives. A tool trained on Tox21 and invitroDB has seen that era’s chemistry and therapeutic classes extensively whether or not the specific compounds appear in its corpus, and these confounds already carry substantial predictive power (Figure 1). Thus, our panel is a harder and better-posed test of generalization to novel compounds, since approval year does not separate its labels as sharply.

**Figure 2:**
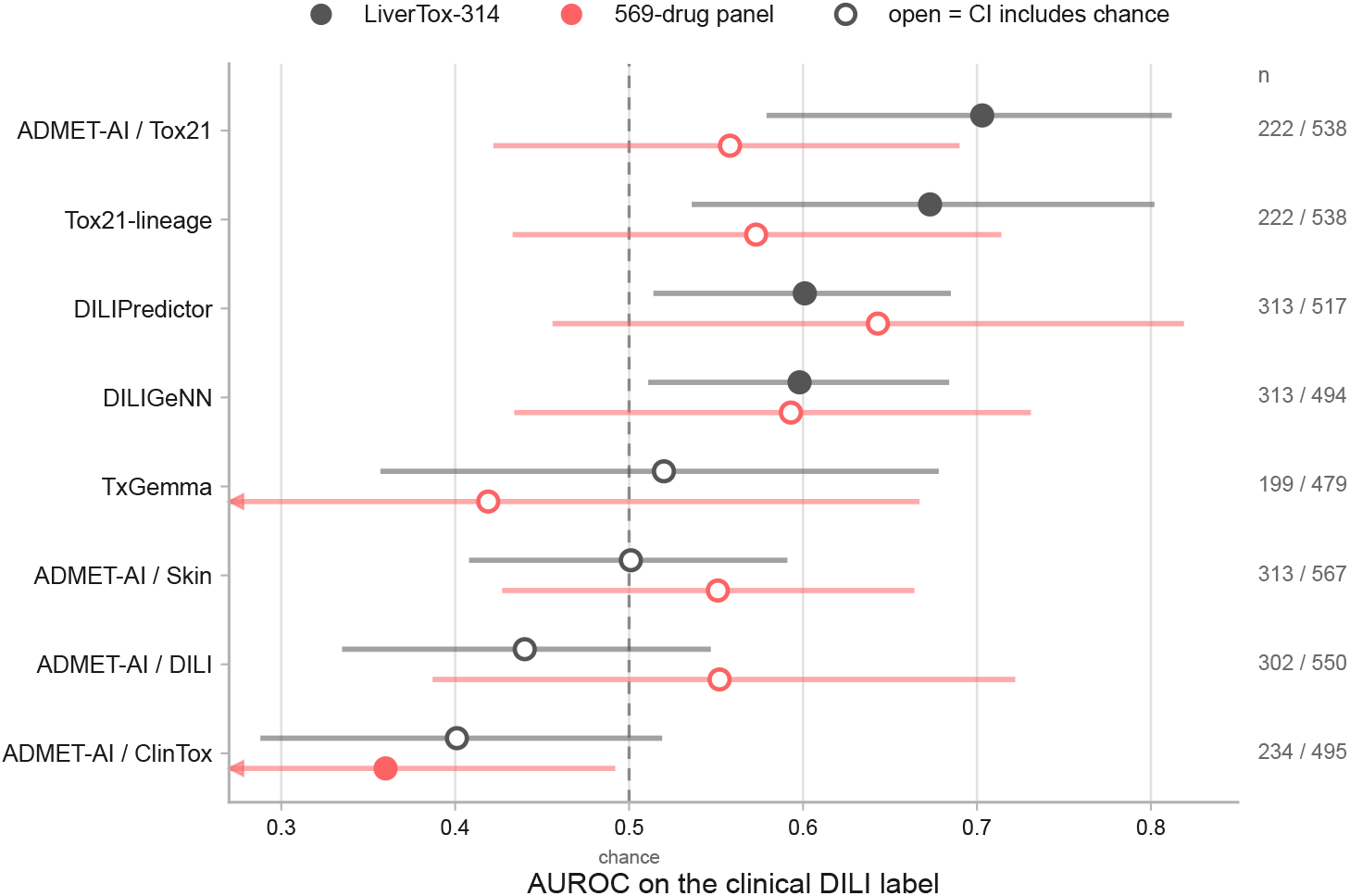
DILI evaluation. Gray: the LiverTox-314 set of Bergen et al.; coral: our 569-drug panel; both scored on DILI labels. Markers are AUROC with 95% bootstrap confidence intervals, open where the interval includes chance (dashed line); arrows mark intervals that extend past the axis. Every value is computed after removing structures matching that tool’s own training set on 2D InChIKey or canonical SMILES, with the number remaining given per row (LiverTox / panel). All tools are frozen.

**Figure 3:**
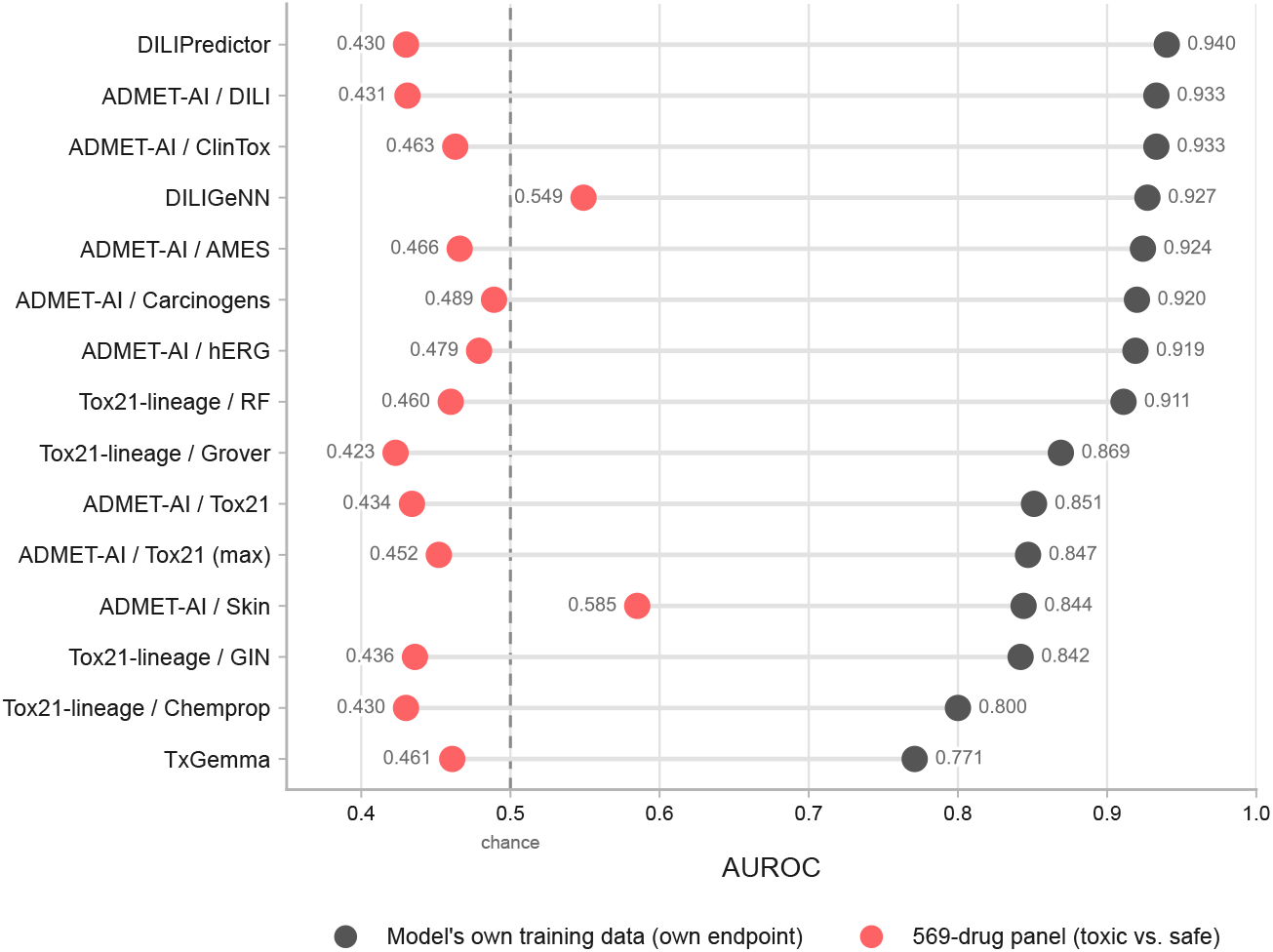
Generalization controls. Each row is one tool. Gray dots show AUROC of each tool on their own dataset. Coral dots show AUROC on the 569-drug panel for the toxic label (black-box warning or withdrawn vs. safe). The dashed line marks chance.

**Figure 4:**
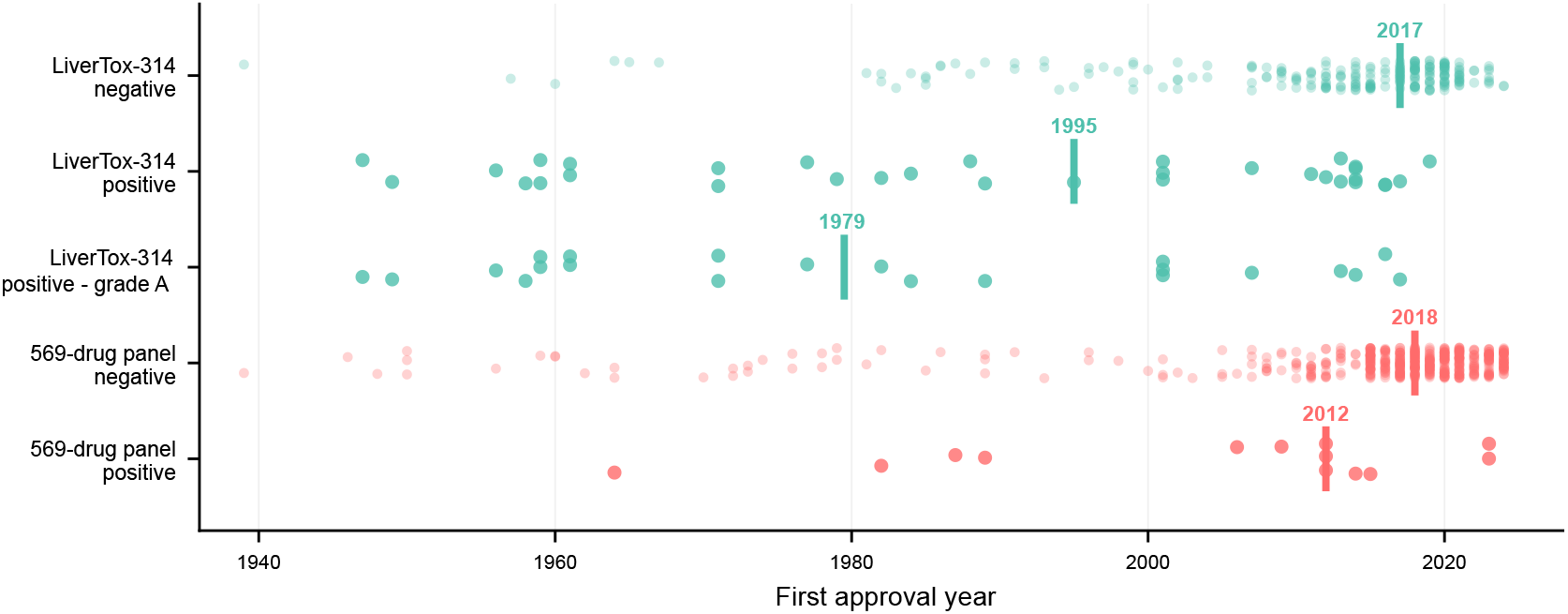
First approval year by dataset and DILI label. Each point is one compound and the vertical bar marks the median. In LiverTox-314 the positives (median 1995) predate the negatives (2017) by 22 years, and the high-confidence grade A positives are older still (1979). Our 569-drug panel narrows that separation to six years (positives 2012, negatives 2018). Both datasets draw their negatives from the same era, so the 22-year gap belongs to LiverTox-314’s positive class alone. Its positives are the textbook hepatotoxins, old enough to sit in almost any training corpus and old enough that approval year by itself predicts the label.

## 5 Conclusions and Limitations

Predicting toxicity before a compound reaches human clinical trials would improve patient safety and spare developers the cost of late-stage failure. Despite the performance reported on their own datasets (Figure 3), available tools cannot reliably predict toxicity on unseen chemistry. No learned molecular representation outperforms a plain fingerprint, so the bottleneck is not the sophistication of the representation. Only one toxicity-specific tool clears chance on our panel, and none matches what approval year alone explains, a predictor containing no chemistry at all. No DILI-specific tool clears chance on our panel, and the external set where some do is one approval year alone largely explains. These conclusions rest on a curated panel that is modest in size, thin for DILI (22 positives), and not balanced enough to rule out every confound, and on regulatory action as a toxicity proxy. A larger panel is future work. Still, a structure encodes what a compound is, while toxicity depends on how much of it reaches which tissue, what it is metabolized into, and in what biological state. Structure encodes none of that. Progress therefore requires measurements that capture exposure and metabolism directly, not better models over structure alone.

## Acknowledgments and Disclosure of Funding

We thank all members of Matterworks, Inc. for their advice and help on completing this work. All authors of this work are employees or shareholders of Matterworks, Inc.

## A Appendix

### A.1 Model classes and evaluation protocols

The paper asks whether molecular structure alone predicts clinical toxicity for drugs unlike those a model has seen. Each Class below is described together with how it informs that question, and the Classes differ in how much each model can learn from the panel (Table 2). The roster covers tools that could be scored on all 569 compounds in batch and whose training compounds were available for a leakage audit, which excludes regulatory QSAR systems such as EPA TEST [Martin, 2020], the OECD QSAR Toolbox [Dimitrov et al., 2016] and Derek Nexus [Lhasa Limited], as well as web servers such as ProTox [Banerjee et al., 2024] and ADMETlab [Xiong et al., 2021]. We say a model is applied *frozen* when no parameter is fit on the panel, and for the Class 2 encoders *frozen* refers to the encoder weights alone, since the linear probe over their embeddings is still fit on panel labels. Every model fit on the panel uses the same 5-fold splits.

#### Class 0: baseline models

Class 0 asks how well simple baselines can do, most of them learning directly from the panel. This shows what can be learned from the panel’s own compounds, as opposed to what must transfer to new chemistry. The Class includes models built on ECFP4 fingerprints [Rogers and Hahn, 2010], on physicochemical descriptors, and on both combined (Table 2), using logistic regression, XGBoost or random forest classifiers. These classifiers are widely used for fingerprint-based prediction models and cover linear, boosted-tree and bagged-tree methods, so our conclusions do not depend on a single choice. It also includes two probes that use no chemistry (approval year and therapeutic class), which show how much of the toxicity label metadata alone can predict. Raw ECFP4 is our representation for in-distribution reference. Models built on it are fit on part of the panel, so any model that beats it must offer something beyond training on compounds that resemble the test compounds. Finally, Class 0 includes a pipeline that predicts bioactivity across 471 invitroDB assays [Dix et al., 2007] from structure and maps it to toxicity. We evaluate it both frozen and with a head fit on panel labels, to separate what the bioactivity layer contributes from what the panel supplies.

#### Class 1: published toxicity tools

Class 1 asks what a drug-development team would get by using published toxicity predictors as released. These tools were trained on their own large corpora, so if structure alone supported toxicity prediction they should carry over to drugs outside those corpora, which is the question we test here. This Class includes the ADMET-AI toxicity heads [Swanson et al., 2024], classifiers from the Tox21 lineage [Mayr et al., 2016], the liver-injury tools DILIPredictor [Seal et al., 2024] and DILIGeNN [Lee and Posma, 2025], and TxGemma [Wang et al., 2025] (Table 2). We test each tool using its released checkpoint, so nothing is fit on our panel. Because a tool may have seen panel compounds in training, we remove from each evaluation the compounds that match that tool’s training set, so that a score reflects performance on compounds it has not seen.

#### Class 2: learned structure encoders

Class 2 asks whether a more expressive representation of structure helps with toxicity prediction. If the limit were the representation, a learned encoder should outperform a fixed fingerprint when both receive the same panel, folds and labels. This Class includes Uni-Mol [Zhou et al., 2023], MolFormer-XL [Ross et al., 2022], ChemBERTa-2 [Ahmad et al., 2022] and Chemprop v2 [Heid et al., 2024], each finetuned on the panel (Table 2). For the three with a pretrained checkpoint we also fit a linear probe on the frozen embeddings, and comparing the probe with the finetuned model separates what the pretrained representation already contains from what adaptation to the panel adds. Because these models are fit on the panel like the Class 0 baselines, they are compared with raw ECFP4, the in-distribution reference, and not with the frozen Classes.

#### Class 3: transfer from animal data

Class 3 asks whether structure–toxicity relationships learned from a different source of labels carry over to the clinical outcomes of the panel’s drugs. If such relationships were general, a model trained on non-clinical data should transfer. This Class applies the Class 0 ECFP4-plus-descriptor recipe, trained instead on 495 ToxRefDB animal-study compounds [Feshuk et al., 2023, Watford et al., 2019] with logistic regression, XGBoost and random forest heads, to the panel without refitting (Table 2). Because the recipe is the same as in Class 0, the training data is the only thing that differs between the two Classes. ToxRefDB is built from guideline animal studies rather than from regulatory action on approved drugs, so its labels differ from the panel’s as well as its chemistry.

### A.2 Sensitivity to the novelty cutoff

To check that the 0.70 cutoff does not drive our results, we scored the frozen tools on a second cohort of 1,945 approved drugs from ChEMBL 37. This cohort has no similarity cutoff and no first-approval cutoff, so its drugs range from 0.06 to 1.00 in similarity to invitroDB. We split it into six similarity bins. Mean AUROC across the 13 readouts stays between 0.52 and 0.60 in every bin and does not systematically increase with similarity (Table 3). Six readouts score slightly higher above the cutoff than below it and seven score lower, and none exceeds 0.61 above it, so a looser cutoff would not change the frozen-tool results. This check concerns similarity to invitroDB and not to each tool’s own training set, which is why, on the panel, we separately remove from each tool’s evaluation the compounds found in that tool’s training set. Similarity is also linked to approval era in this cohort (*r* = *−*0.51).

### A.3 Whether the below-chance scores reflect a failure to generalize

Every frozen tool that could be scored clears chance decisively on its own training data, from 0.771 to 0.940, against 0.43 to 0.59 for the same models on the panel. DILIPredictor reaches 0.940 on its own training set and 0.430 on the panel, and the ADMET-AI DILI head 0.933 against 0.431. Every model whose significance we report is covered in Figure 3.

